# One-Class Bioacoustic Detector for Monitoring the Critically Endangered Pied Tamarin (*Saguinus bicolor*)

**DOI:** 10.1101/2025.10.11.681843

**Authors:** Juan G. Colonna, Tainara V. Sobroza, Marcelo Gordo, Eduardo F. Nakamura, Alejandro C. Frery

## Abstract

The pied tamarin (*Saguinus bicolor*) is a critically endangered primate with a small geographic range that includes fragmented urban forest mosaics in Amazonia, where habitat subdivision and anthropogenic actions complicate its survival and monitoring. Passive acoustic monitoring (PAM) offers a convenient, noninvasive way to track this species, yet open-set rainforest soundscapes make single-species detection challenging. We present a machine-learning pipeline with a very low false-positive rate, appropriate for downstream inference. The method combines a band-pass filter (5 kHz to 10 kHz), Perch bioacoustic embeddings (deep learning), and a One-Class SVM (OCSVM) applied to sliding windows of continuous audio recordings to detect *S. bicolor* calls. We train on a reduced dataset of labeled calls and validate against diverse out-of-class audio (birds, anurans, anthropophony, and geophony/insects), then test on long, cross-site recordings. The approach achieves high discrimination on held-out negatives and produces very low false-positive rate in continuous, real-world audio, with a precision of 0.86. Finally, we pair detections with a single-site occupancy model in a cross-site setting to illustrate end-to-end utility for conservation monitoring and to estimate the false-negative detection probability in recordings from pied tamarin populations in a different geographic region. Our strategy provides a tool for PAM of *S. bicolor* that requires minimal manual labeling effort and can be adapted to other open-set, single-species monitoring scenarios. We grant reproducibility by releasing a Python package (`sauim-detector`), installable via `pip`, that processes an audio file and produces detection timestamps as an Audacity label file (`txt`), enabling faster manual verification.

## Introduction

The pied tamarin (*Saguinus bicolor*, Figure 1a) is a small primate of the Brazilian Amazon. This species is endemic to a geographically restricted region of approximately 8354 km^2^ (Lagroteria et al., 2024; Albernaz et al., 2026). Unfortunately, this geographic range includes the city of Manaus, the second-largest urban agglomeration in the Brazilian Amazon. Due to its sensitivity to habitat loss and fragmentation, and its importance to the Amazonian environment through seed dispersal (Fernandes et al., 2024), the species has been monitored for more than four decades (Ayres et al., 1982).

**Figure 1.**
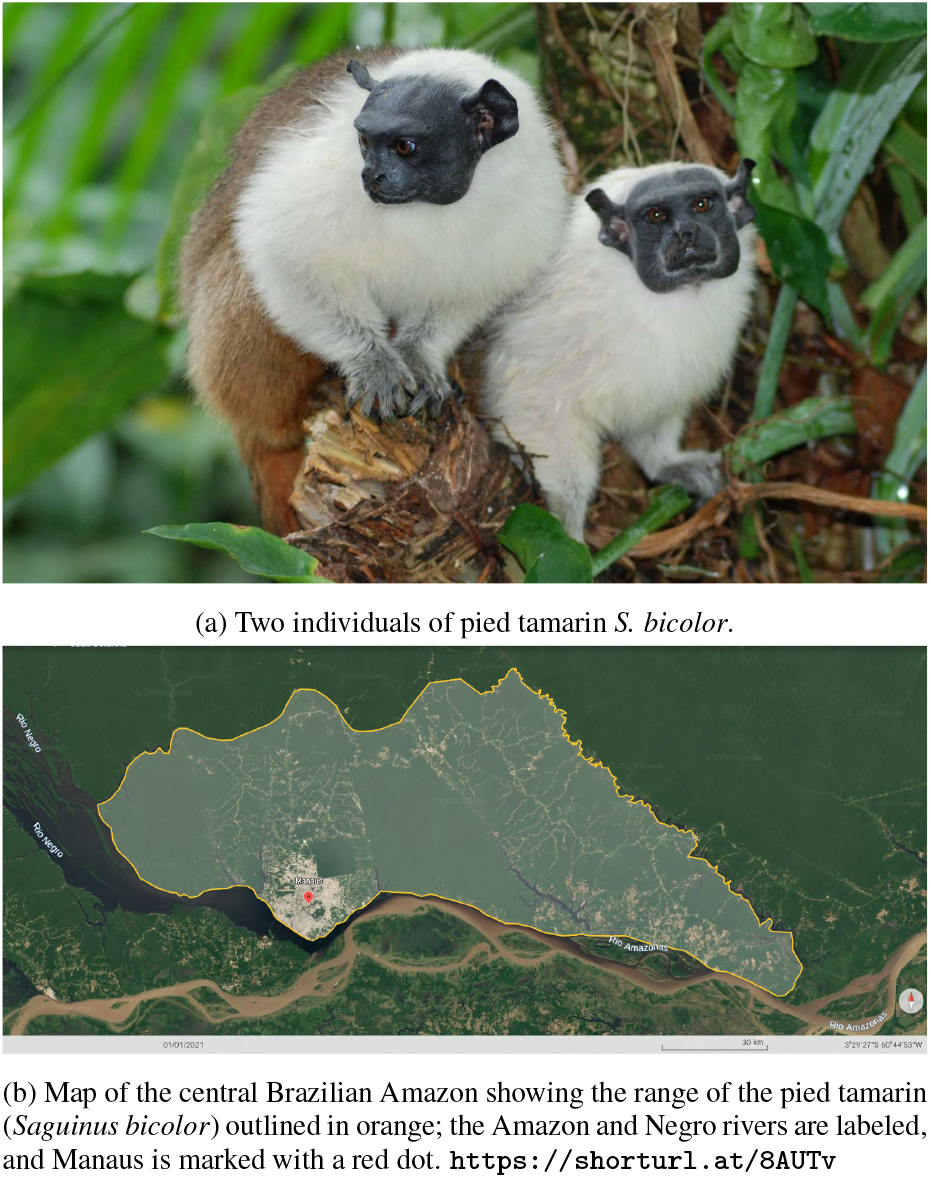
(a) Pied tamarins (*S. bicolor*) photographed in an Environmental Protection Area (APA) on the Federal University of Amazonas campus. (b) The species’ geographic range overlapping with the city of Manaus in the Brazilian Amazon. Adapted from Lagroteria et al. (2024); Albernaz et al. (2026).

The *S. bicolor* is listed on the IUCN Red List of Threatened Species as a critically endangered species, with its assessment updated in 2021 (Gordo et al., 2021). Within the city of Manaus, several urban forest protected areas have served as refuges for the pied tamarin; however, because these are fragmented urban forests, survival remains challenging (Vidal and Cintra, 2006). Vehicle collisions, electrocution on power lines, and habitat degradation due to human encroachment are among the main threats (Gordo et al., 2013). In 2011, the Brazilian government created a National Plan for the conservation of the *Sauim-de-coleira*. Initiatives have been coordinated to study and monitor populations to support evidence-based management with the aim of safeguarding the species (ICMBio, 2011).

Among the several available ways to detect the presence of the *S. bicolor*, passive acoustic monitoring (PAM) systems are among the least intrusive methods (Sobroza et al., 2025). With PAM, autonomous acoustic recorders capture the Amazonian soundscape, and the presence of the pied tamarin can be subsequently inferred from its vocalizations (calls). Recorders offer wide detection ranges and approximately 360^°^ coverage without requiring line of sight, as would a camera trap (Crunchant et al., 2020). However, the abundance of non-target sounds and the dynamic forest background impose a substantial challenge for reliable acoustic detection.

High-quality acoustic classifiers for other animal groups have been proposed (e.g., BirdNET for birds (Kahl et al., 2021) and howler monkeys (Wood et al., 2023a); anuran call detectors (Colonna et al., 2016)), but none specifically target *S. bicolor*. Training a binary or multi-class classifier for a single target species poses a fundamental challenge: how can we adequately represent the negative class when the acoustic environment contains hundreds of potential non-target species, plus anthropogenic and environmental sounds? The Amazonian soundscape is rich, including birds, anurans, other mammals, insects, geophonies (e.g., wind, rain, rivers) (Xavier et al., 2024), and anthropophonies (e.g., traffic, voices, machinery) due to proximity to urban areas or human incursions (Sobroza et al., 2024). This diversity, and the fact that the set of non-target sounds is effectively unbounded and shifts across sites and seasons, characterizes an open-set recognition problem (Geng et al., 2021). Consequently, a one-class or anomaly-detection approach is appropriate for *S. bicolor* detection, as we adopt in this work.

Fortunately, the pied tamarin has a distinctive, nearly unique call signature, except in zones of sympatry where golden-handed tamarins (*S. midas*) may shift their calls toward the pied tamarin’s pattern (Sobroza et al., 2021). The characteristics of *S. bicolor* vocalizations can be seen in the spectrogram in Figure 2b. A spectrogram is a two-dimensional representation of an acoustic signal, with time on the horizontal axis, frequency on the vertical axis, and energy encoded as color intensity. This representation enables detailed analysis of the temporal and spectral structure of animal vocalizations and is essential in acoustic studies.

**Figure 2.**
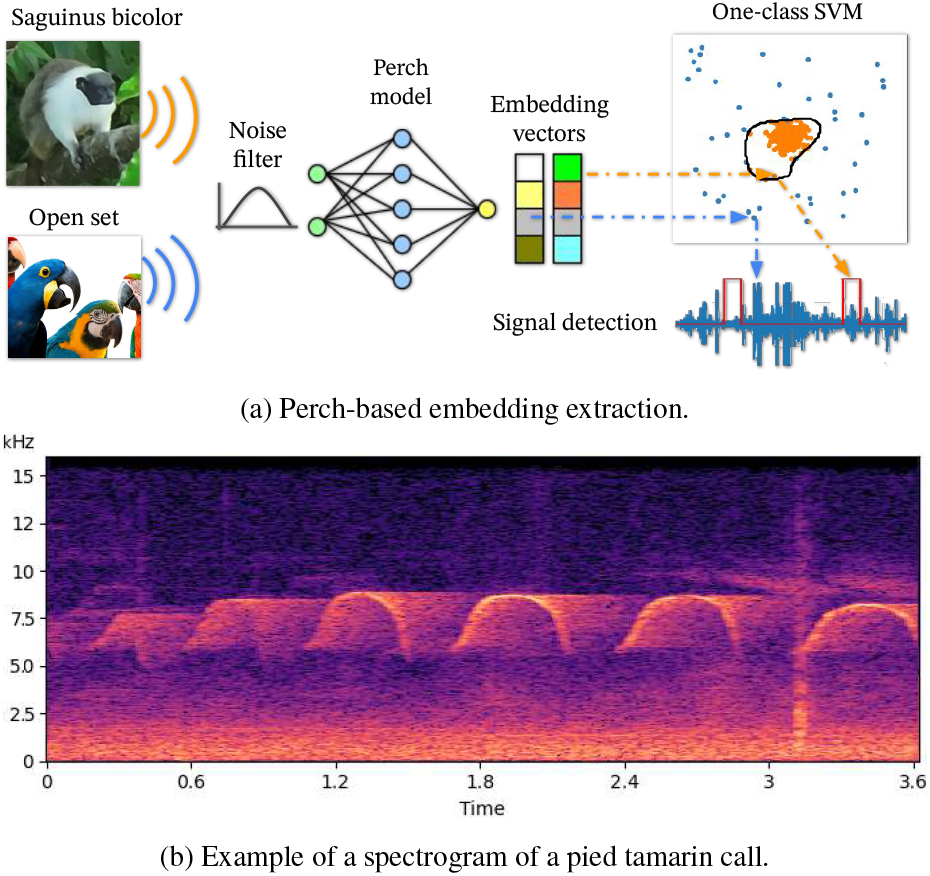
(a) Conceptual overview of the proposed approach. (b) A spectrogram showing a typical call of this species.

In *S. bicolor*, call energy is concentrated predominantly between approximately 6 kHz to 9 kHz. The calls consist of a sequence of arch-shaped patterns (i.e., syllables) repeated over the course of the signal, typically comprising one to six syllables with a total duration of roughly 2 s to 5 s (Sobroza et al., 2017). Such variation may reflect different behavioral contexts, including group cohesion, territory defense, or even adjustments to anthropogenic sounds (Sobroza et al., 2024). This domain knowledge is incorporated into our method via a band-pass filter that targets this frequency band (Section 3.5).

The distinctive and consistent acoustic features of pied tamarins (e.g., call duration, syllable repetition rate, and characteristic pattern shapes) provide reliable species-specific signatures that can be leveraged in passive acoustic monitoring (PAM) systems, enabling non-invasive detection and long-term population tracking. Because our objective is to autonomously and specifically monitor a single species of interest, an open-set recognition formulation is required to handle signals from previously unseen classes (Section 2).

In the present study, *S. bicolor* vocalizations constitute the sole “known” class, and a One-Class Support Vector Machine (OCSVM) is applied to detect calls consistent with this class while rejecting sounds from other species and background noise. Conceptually, one-class classification can be viewed as a special case of open-set classification, in which only a single target class is modeled during training (Khan and Madden, 2014). In our method, an incoming audio recording is first normalized to ±1 amplitude, band-pass filtered (e.g., 5 kHz to 10 kHz), and segmented into five-second windows. Then each segment is passed through the Perch bioacoustic embedding model to obtain an embedding vector. Finally, these vectors are classified by an OCSVM trained only on pied tamarin calls (Figure 2a).

While open-set classification assigns inputs to one of several known classes or rejects them as unknown, one-class classification defines a boundary that encloses only the target class in feature space (cf. Figure 2a), treating all other inputs as unknown or anomalous (Marques et al., 2023). Therefore, our approach addresses the open-domain challenge of bioacoustic monitoring in complex soundscapes, where signals from unobserved classes are inevitably present.

## 2. Problem formulation

An open-set problem is a setting in which, at test time (deployment), classes not seen during training may appear (Scheirer et al., 2013). The model must both recognize the known classes and be able to reject samples from unknown classes (rather than forcing a misclassification into a known class).

Formally, let *K* be the set of classes available during training and *C* the set of classes that may appear at test time, with *K* ⊆ *C*. The {unknown} classes are *U* = *C* \ *K*. An open-set classifier implements a decision rule *f* : X → *K* ∪ unknown that (i) assigns correct labels to inputs *x* ∈ X from classes in *K*, and (ii) rejects inputs from *U*, avoiding high-confidence predictions in regions of the feature space without training support. In our case, the cardinality of *K* is one (|*K*| = 1); therefore, we adopt a one-class model based on the One-Class Support Vector Machine (Schölkopf et al., 2001).

In this study, each *Saguinus bicolor* vocalization is represented by a 1280-dimensional embedding vector extracted from a 5 s audio segment using Google’s Perch bioacoustic model (Google Research, 2025). Formally, the feature extractor can be seen as a mapping *f* : *X* → *V, x* ↦ **v** ∈ ℝ^1280^, where *X* denotes the space of 5 s acoustic waveforms and *V* the resulting feature space. We hypothesize that these embedding vectors capture salient spectral-temporal patterns of the calls, enabling robust separation of *S. bicolor* vocalizations from all other sounds in a complex and diverse rainforest soundscape.

To model the target class, we apply OCSVM, which seeks to find a decision boundary in the highdimensional embedding space that encloses most of the training vectors while allowing a small fraction of them to fall outside (Schölkopf et al., 1999). In its primal form, the OCSVM solves:

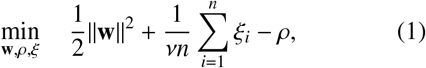

subject to (**w** · **v**_*i*_) ≥ ρ − ξ_*i*_, ξ_*i*_ ≥ 0, *i* = 1, …, *n*, where:

- **v**_*i*_ ∈ ℝ^1280^ is the embedding vector of the *i*-th vocalization segment;
- **w** is the normal vector defining the separating hyperplane in the feature space;
- ρ is the offset defining the decision threshold;
- ξ_*i*_ are slack variables that permit some training points to be considered outliers;
- *n* is the number of training examples;
- ν ∈ (0, 1] controls the maximum fraction of training outliers and the minimum fraction of support vectors.

The decision function for a new embedding vector **v** is *f* (**v**) = sign (**w** · **v** − ρ), returning +1 if the segment is classified as a *S. bicolor* vocalization and −1 if classified as an anomaly (i.e., another species, environmental sound, anthropogenic noise, etc).

In the OCSVM formulation (Eq. 1), the hyperparameter ν ∈ (0, 1] plays a dual role. First, ν sets an upper bound on the fraction of training samples allowed to lie outside the learned decision boundary. Second, it sets a lower bound on the fraction of support vectors and, together with the kernel parameters (e.g., γ for the RBF kernel), governs the complexity of the decision function. In the context of *S. bicolor* acoustic monitoring, setting ν too high yields a very tight boundary that may overfit the variability of training calls, misclassifying legitimate vocalizations as anomalies. Conversely, setting ν too low produces a looser boundary that risks admitting non-target sounds, e.g., other animal vocalizations, geophonic events (e.g., wind, rain), or anthrophonic noise, as *S. bicolor* calls.

## 3. Materials and method

We used five datasets in the experiments that follow. The main dataset contains recordings of *S. bicolor* vocalizations captured outdoors under real Amazonian rainforest conditions. The recordings were made using a Sennheiser ME67 unidirectional microphone matched to a Zoon H4N recorder. The remaining four datasets comprise bird and anuran calls, as well as anthropophony, geophony, and insects noise. This diversity of sounds is intended to cover scenarios likely to occur during field monitoring with acoustic sensors.

### 3.1. S. bicolor and background noises dataset

The *S. bicolor* dataset consists of two sets of recordings: 32 recordings from Bosque da Ciência at the Instituto Nacional de Pesquisas da Amazônia (INPA), and 38 recordings from Mindú Park. The INPA recordings range from 2 to 13 minutes, totaling approximately 2 hours and 17 minutes, while the Mindú recordings range from 33 seconds to 27 minutes, totaling approximately 2 hours and 38 minutes. These sets come from two different groups of individuals and exhibit distinct background noise conditions. Having multiple specimens is desirable for testing the transferability and generalization of the trained model, as calls may share global acoustic characteristics while also showing subtle, individual-specific differences.

From the first group, the recordings were segmented into five-second windows whenever a pied tamarin vocalization was present, resulting in 118 five-second segments containing only pied tamarin vocalizations. As the vocalizations are sparse throughout the recordings, this procedure recovered only the necessary segments for training, which together resulted in approximately 10 minutes of audio.

We also extracted 179 randomly selected segments containing only background noise. These noise segments were merged with the 118 call segments, totalizing 297 feature vectors composing a binary dataset. This set was split (holdout) using the 70–30% rule, resulting in 82-36 calls for training and testing as the target class, and 125-54 samples for training and testing as the non-target class. Thus, the OCSVM was trained using only the target class with 82 *S. bicolor* calls. Each sample was labeled by the OCSVM as an inlier (target) or outlier (non-target). We refer to this setup as the “Background” experiment in our figures and tables.

### 3.2. Birds and Anurans datasets

Common sources of bioacoustic events in rainforests are vocalizations produced by birds and anurans (frogs and toads). Many bird species vocalize in high-frequency bands, increasing the likelihood of overlap with *S. bicolor* calls and, consequently, leading to misclassification. In this study, we curated 30 recordings from 26 bird species obtained from the collaborative Xeno-canto repository^1^, totaling 130 five-second record segments. We selected only recordings from the city of Manaus and its surroundings, overlapping the geographic range of the *S. bicolor* individuals used to train our OCSVM.

The anuran dataset includes 60 recordings from ten species, some of which occur frequently in the same areas as *S. bicolor*, while others are more broadly distributed across the Amazon. In total, we obtained 122 five-second acoustic segments. Note that most of these amphibians, which could potentially confuse the machine learning algorithm, vocalize at night, whereas the pied tamarin vocalizes during the day. Complete lists of bird and anuran species are provided in Section Appendix A. We refer to these two datasets as “Birds” and “Anurans”; they serve as outlier sets to evaluate our method’s performance on acoustic events outside the target class.

### 3.3. Anthropophony dataset

The anthropophony (human-generated sounds) dataset was assembled from the ESC-50 corpus (Piczak, 2015). We concatenated variable-length segments into a single long recording comprising three airplane segments, one car horn, three city traffic segments (including horns), three chainsaw segments, two church bells, one applause segment, three coughs, four miscellaneous engine sounds (e.g., tractor, motor), three fireworks, three hand saws, two helicopters, three trains, and three different siren recordings.

These sources reflect plausible human activities near protected areas—for instance, fireworks during soccer match days can be audible across urban reserves; engines and motors are common at reserve edges; chainsaw and hand saw sounds may indicate illegal logging; and helicopters occasionally overfly monitored sites. Although some of these sounds (e.g., bells or trains) are not typical of the areas occupied by the pied tamarin, these negative samples help assess our method’s capability to generalize to other scenarios. We standardized these clips into 84 five-second segments, which we use as the “Anthropophony” set.

### 3.4. Geophony and insect sounds

The geophony and insect dataset collected from SoundCloud includes three crackling fire segments, three cricket segments, four additional insect recordings (mostly flies), five rain segments, three sea wave segments, four thunderstorm segments, two water drop segments, and five wind segments. Although some of these sounds, for example sea waves, are not common in Amazonia, they add diversity to the negative class in this experiment. Because our strictly biophonic material was limited, we grouped common insect sounds (e.g., flies, crickets, bees) with geophony for this class, as both can introduce broadband or quasi-stationary energy that may mask target calls. After standardization, this collection totals 77 five-second segments, which we refer to as the “Geophony” set.

### 3.5. Band-pass noise filter

Figure 2b shows that the energy of *S. bicolor* vocalizations is concentrated in the 5 kHz to 10 kHz band, whereas rainforest sounds lie outside this range or occasionally overlap it. For example, in the 3.1 s to 3.6 s interval, this spectrogram exhibits transient events that partially overlap the calls. Such overlapping interference degrades the quality of features extracted by the Perch model and can reduce detection/recognition performance. To mitigate this, we include a Butterworth band-pass filter centered on the target band as the first stage of our classification pipeline. Butterworth lowor high-pass filters are widely used in bioacoustics for their simple parameterization and smooth, monotonic magnitude response (Miksis-Olds and Tyack, 2009; Laidre et al., 2024).

We applied a fourth-order (*N* = 4) IIR Butterworth band-pass filter with cutoff frequencies *f*_*c*_ ∈ [5, 10] kHz, implemented as cascaded second-order sections (SOS). The Butterworth response is maximal and flat in the passband and monotonic across the spectrum, with an asymptotic roll-off of approximately 6*N* = 24 dB/octave on each side. Figure 3 shows the magnitude response (dB) for a sampling rate *f*_*s*_ = 32 kHz. Therefore, our recorded signals were processed with zero-phase forward–backward filtering using `sosfiltfilt`^2^ from the SciPy Python library, which eliminates phase distortion while preserving amplitude characteristics. The effect of this filter on *S. bicolor* vocalizations is discussed in Section 6.

**Figure 3.**
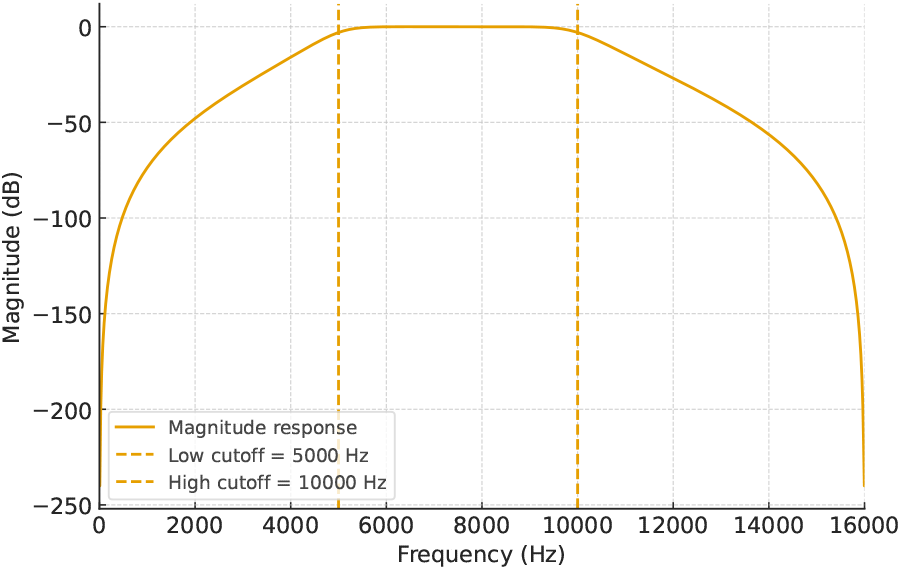
butterworth band-pass filter.

### 3.6. Perch: bioacoustic embedding model

We used embeddings from the Perch bioacoustic model available in the Kaggle public repository (Google Research, 2025), which is based on an EfficientNet-B1 Convolutional Neural Network (CNN) architecture with approximately 7.8 M parameters trained on a global corpus of bird vocalizations (Xeno-Canto) with more than 15 k species (Ghani et al., 2023). This model expects input as single-channel five-second audio segments sampled at 32 kHz; it outputs logits and an embedding vector for each segment.

The Perch model has been compared extensively with BirdNET, AudioMAE, YAMNet, and VGGish, showing strong transfer-learning performance across diverse bioacoustic tasks (Ghani et al., 2023). Derived models, such as SurfPerch, further demonstrate adaptability to non-avian domains (e.g., tropical reefs) (Williams et al., 2025). Importantly, Perch embeddings are highly transferable and data-efficient, enabling few-shot training and an active-learning loop that rapidly surfaces additional samples via vector search, yielding high-quality detectors with modest annotation effort (Dumoulin et al., 2025). These properties make Perch an ideal choice for our approach given our small annotated dataset. Here, we further show that Perch transfers to primates by training a one-class SVM on its embeddings to perform open-set detection.

## 4. Experimental setup

In this study, we adjusted ν (Eq. (1)) through a grid search over [0.01, 0.10] using Leave-One-Out CrossValidation (LOOCV), selecting the value that maximized the area under the ROC curve (AUC) on the described held-out training set (Section 3.1). This optimization was repeated both with and without the band-pass filter. This criterion yielded a balance between sensitivity to target vocalizations and robustness to the heterogeneous Amazonian soundscape. The kernel function was chosen as the radial basis function (RBF), which is well-suited for modeling non-linear class boundaries in high-dimensional spaces, and the γ hyperparameter was automatically set according to the rule 1/features, corresponding to the inverse of the 1280dimensional embedding size. The OCSVM implementation was provided by the `scikit-learn` Python library (Pedregosa et al., 2011).

All codes to reproduce our results are provided as ready-to-run, interactive Jupyter notebooks, together with the data used in our experiments, at https://github.com/juancolonna/Sauim/tree/main.

## 5. Results

Our experimental results are divided into three main sections. The first section addresses the classification problem using the five datasets described in Section 3. The second addresses detection in long recordings (Sec. 6). Section 7 shows how a bioacoustic classifier can be coupled with an occupancy model to estimate true- and false-positive detection probabilities (*p*_11_ and *p*_10_) conditional on site occupancy. We conducted classification and detection experiments with and without a band-pass filter to assess its impact on the method’s performance.

### 5.1. Classification

Our method comprises a five-stage pipeline: (i) load each recording and resample to the Perch input rate (*f*_*s*_ = 32 kHz); (ii) apply the Butterworth band-pass filter; (iii) segment the audio with a five-second sliding window; (iv) pass all segments through the Perch neural network to obtain embedding (feature) vectors; and (v) train the OCSVM on the resulting embeddings.

Of the 118 five-second *S. bicolor* call segments, 82 were used to fit the OCSVM (target-only training) using LOOCV. The remaining 36 call segments were combined with 54 background-noise segments to form the test set. Figure 4 compares confusion matrices obtained with and without band-pass filtering prior to feature extraction.

**Figure 4.**
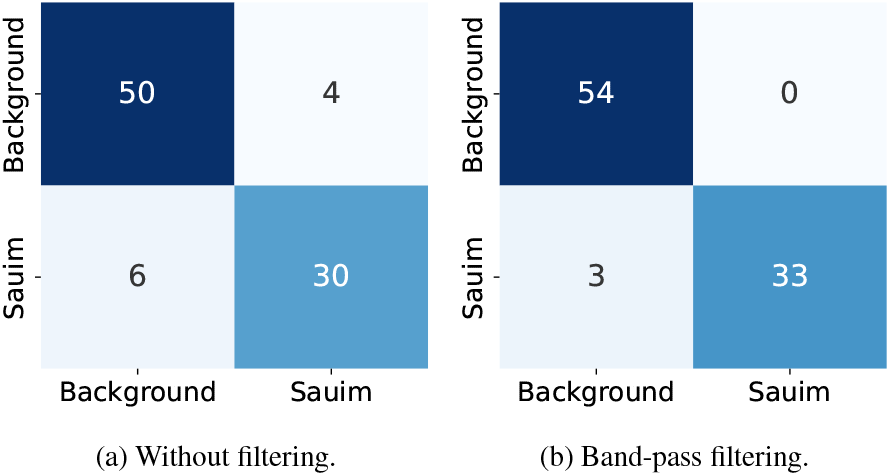
Confusion matrices comparing call detection before and after band-pass filtering on the test set. Rows correspond to true labels; columns to predicted labels. The band-pass filtering eliminates few false positives (4 → 0) and false negatives (6 → 3).

The OCSVM outputs a decision score *s*, with *s* > 0 indicating inliers (call segments) and *s* < 0 indicating outliers (non-target). Segments lying on the soft-margin decision boundary are assigned the label 1. We adopt the rule 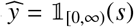, i.e., classify as inlier when *s* ≥ 0. The band-pass filter eliminated the four false positives that appeared when the recordings are not filtered (Figure 4b). Additional confusion matrices for the remaining negative classes (anurans, birds, anthropophony, and geophony+insects) are shown in Figure B.10 provided in Appendix B.

Overall, there is a trade-off between the false-positive rate (FPR) and the true-positive rate (TPR or sensitivity) when varying the decision threshold over the OCSVM score *s* (Figure 5). Each Receiver Operating Characteristic (ROC) curve is obtained by classifying a segment as target whenever *s* ≥ τ and sweeping the threshold τ across the score range. Curves closer to the upper-left corner indicate better performance, whereas the diagonal corresponds to random guessing (Fawcett, 2006).

**Figure 5.**
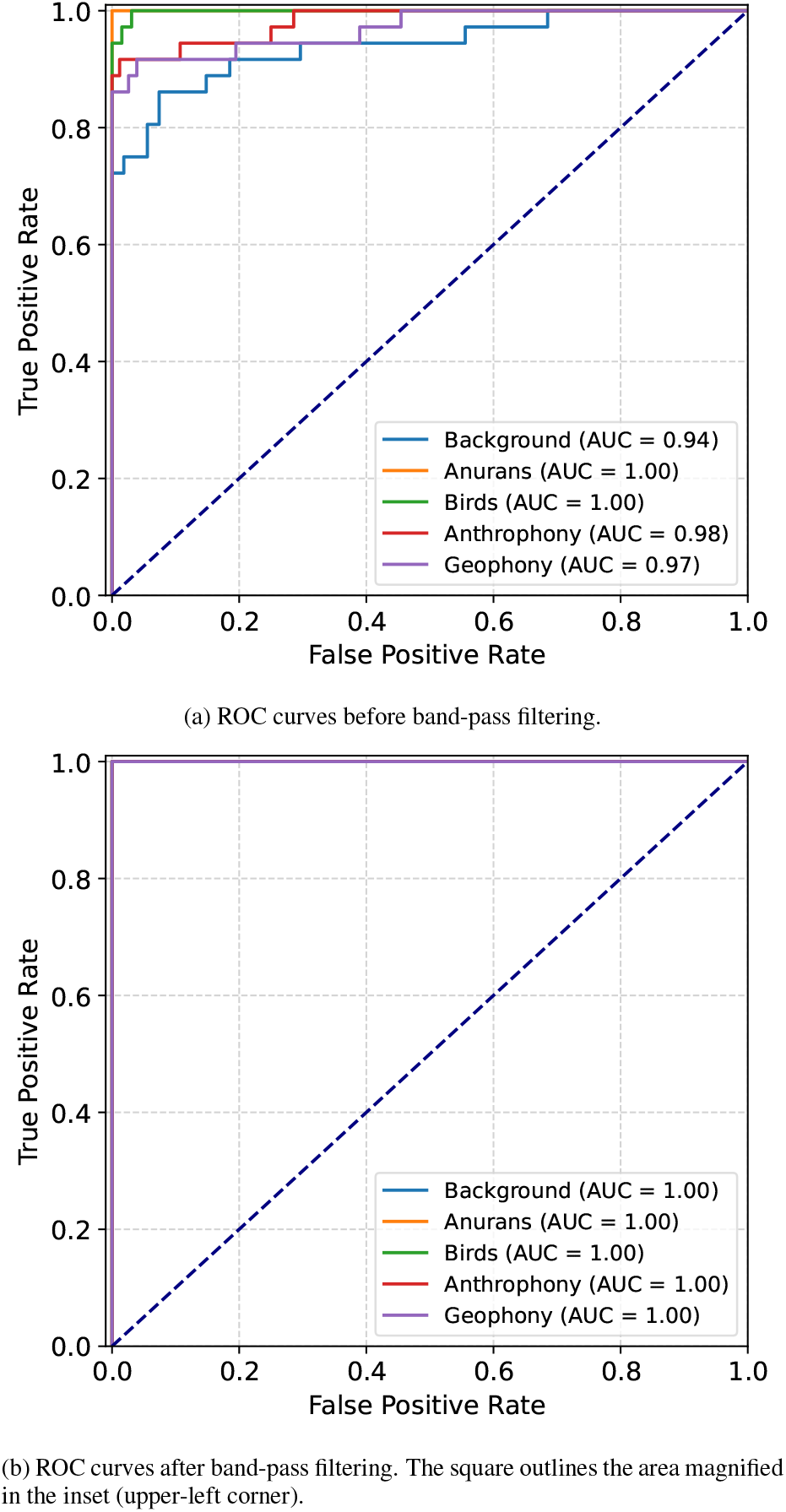
Comparison of ROC curves for OCSVM detection before and after band-pass filtering; most curves overlap, with filtering yielding a slight up-and-left shift (higher AUC).

The area under the ROC curve (AUC) summarizes performance across all thresholds. The AUC ranges from 0 to 1, with 0.5 indicating chance level and 1.0 a perfect classifier (Fawcett, 2006). Probabilistically, AUC = Pr (*s*^+^ > *s*^−^), i.e., the AUC is the probability that a randomly chosen a target call receives a higher score than a randomly chosen a non-target call. Because the AUC is threshold-independent and invariant to any monotonic transformation of *s*, it provides a robust, single-number measure of separability for the OCSVM in this setting. It is evident that applying the band-pass filter prior to feature extraction and classification improves AUC and makes the classifier less sensitive to variations in the decision threshold τ (Figure 5b). In other words, the classes become better separated.

To build intuition about this separability, we performed dimensionality reduction using Principal Component Analysis (PCA) to project the 1,280 dimensional embedding vectors onto two dimensions. *S. bicolor* call segments are well separated in the feature space, facilitating OCSVM classification (Figure 6a). Zooming in on the target class, it highlights which of the samples were marked as outliers during OCSVM training (Figure 6b). Beyond PCA, other analyses can be performed to evaluate the quality of the embeddings. For instance, compactness (within-group cohesion) of the target-class vectors are evident when they are clustered using cosine similarity (Figure D.12 in Appendix D). However, a deeper analysis of embedding quality is beyond the scope of this work.

**Figure 6.**
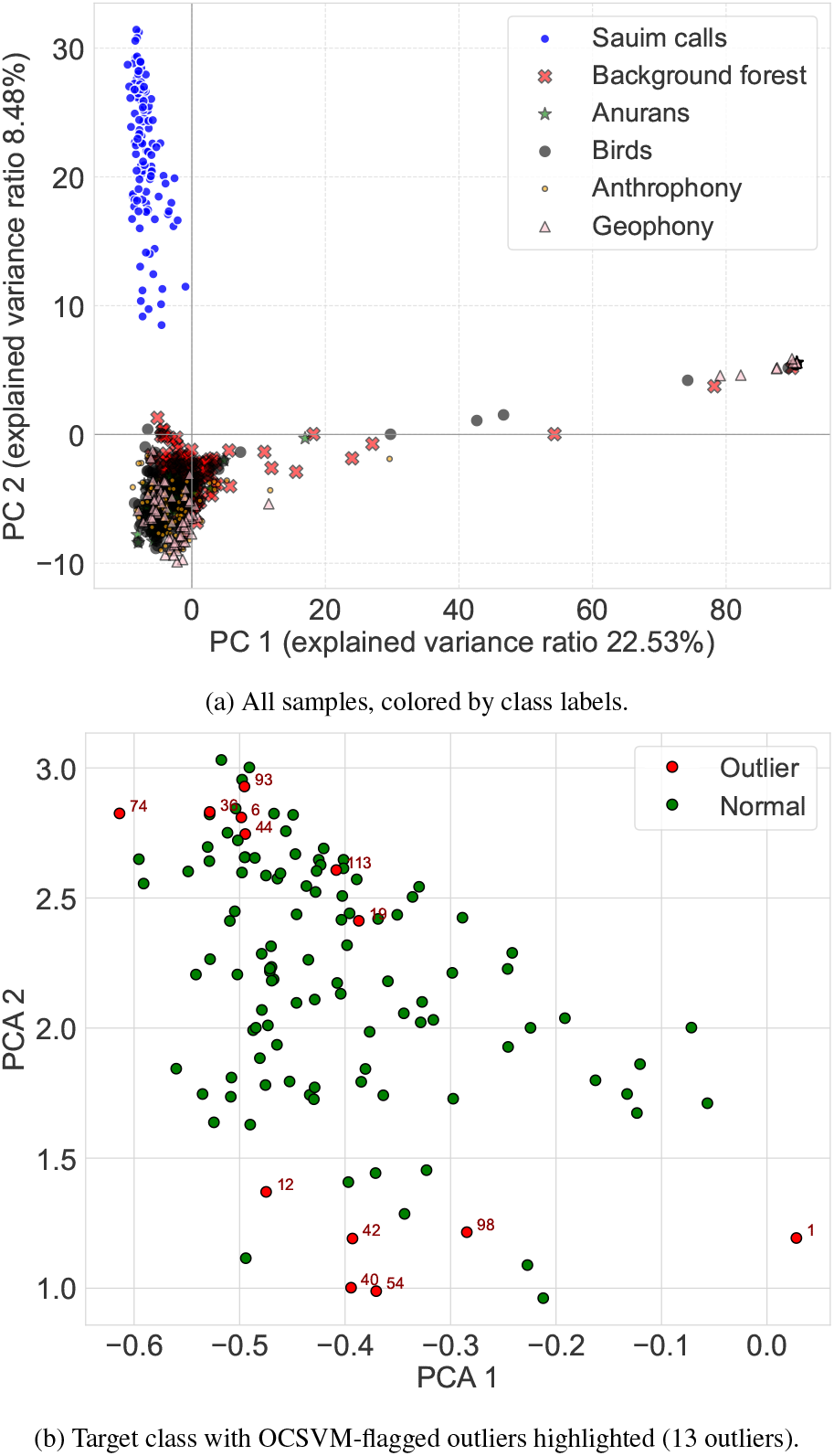
Two-dimensional projection of call-segment embeddings onto the first two principal components (PCA). Panels: (a) all classes;

Finally, Table 1 shows the performance of our OCSVM across five metrics: accuracy (Acc), precision (Pre), recall (Rec), weighted F-score (F1), and AUC—evaluated against the five datasets used as negative classes, yielding an open-set scenario. All these metrics were computed on the band-pass–filtered signals (Table 1). The weighted F1 score is computed as

**Table 1.**
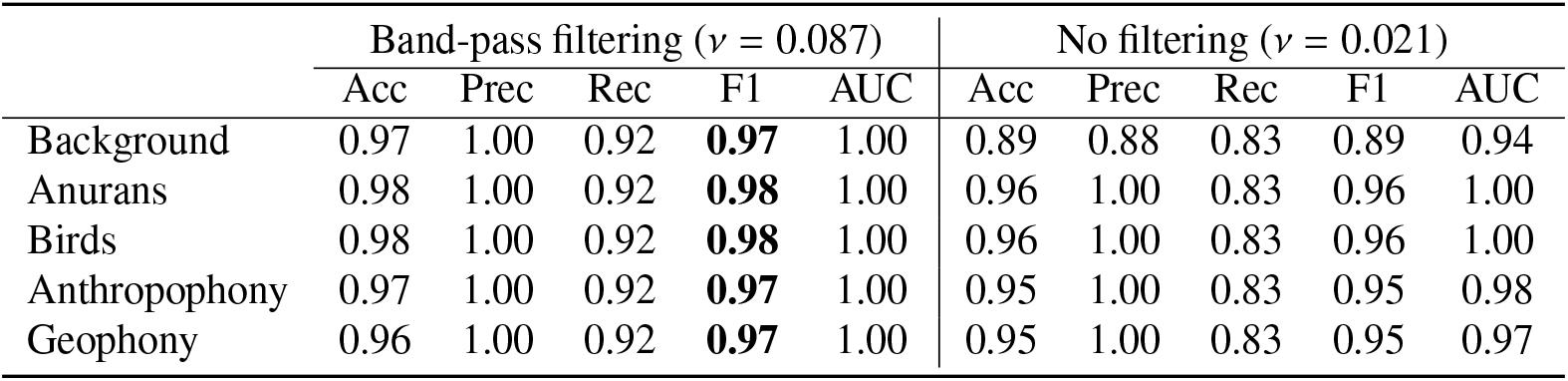
Improvements after applying the band-pass filter. The best F1 scores are highlighted in bold. The optimal ν value found by LOOCV is presented for each case.

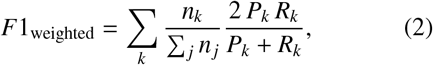

where *k* indexes the classes (here, inlier vs. outlier), *n*_*k*_ is the support (number of true samples) of class *k, P*_*k*_ = TP_*k*_/TP_*k*_+FP_*k*_ is precision, and *R*_*k*_ = TP_*k*_/TP_*k*_+FN_*k*_ is recall for class *k*.

The F1 remains above 97 % in all cases, with only minor variation across datasets (Table 1). This stability reflects (i) the strong separability induced by the Filter+Perch+OCSVM pipeline (consistent with the overlapping ROC curves and high AUC), and (ii) the weighting by class support in the F-score, which reduces sensitivity to small fluctuations in the minority class. Overall, the results indicate excellent performance of the proposed pipeline for *S. bicolor* call detection under simulated open-set conditions.

However, a word of caution is worth mentioning: the held-out dataset is small compared to the training set size. Although we adopted a standard evaluation procedure commonly used in machine learning, the scarcity of *S. bicolor* data makes fair and objective evaluation challenging, occasionally producing near-perfect values for some metrics. To mitigate this limitation and improve the robustness of the evaluation, the next two sections assess the generalization capability of the model using data from another pied tamarin community in a cross-site configuration.

## 6. Use case: a detection example

In our pipeline, ‘classification’ and ‘detection’ are tightly coupled: to detect the target species we classify short segments extracted from a long recording. The experiment in this section simulates scanning a long audio record file to detect *S. bicolor* calls. A real deployment requires the classifier to perform well on recordings from new geographic sites not present in training. Although we search for the same species, the forest background and co-occurring fauna can differ substantially across sites, producing a distribution shift in the embeddings/features and thus testing the generalization of our method.

For this detection example, we used a ten-minute-long recording from Mindú Park for validation. Note that Mindú Park is geographically distinct from Bosque da Ciência, and the *S. bicolor* groups at the two sites are not in contact. Surroundings also differ, leading to different interference sources in the audio. The recording was human-annotated: the onset of each *S. bicolor* vocalization was time-stamped (Figure 7). We processed the waveform (amplitude normalized to [−1, 1]) with a five-second sliding window and a one-second hop (80 % overlap), yielding 559 segments (and therefore 559 embedding vectors). The OCSVM then labeled each segment as inlier (sauim) or outlier (non-sauim). In total, 51 detections were registered.

**Figure 7.**
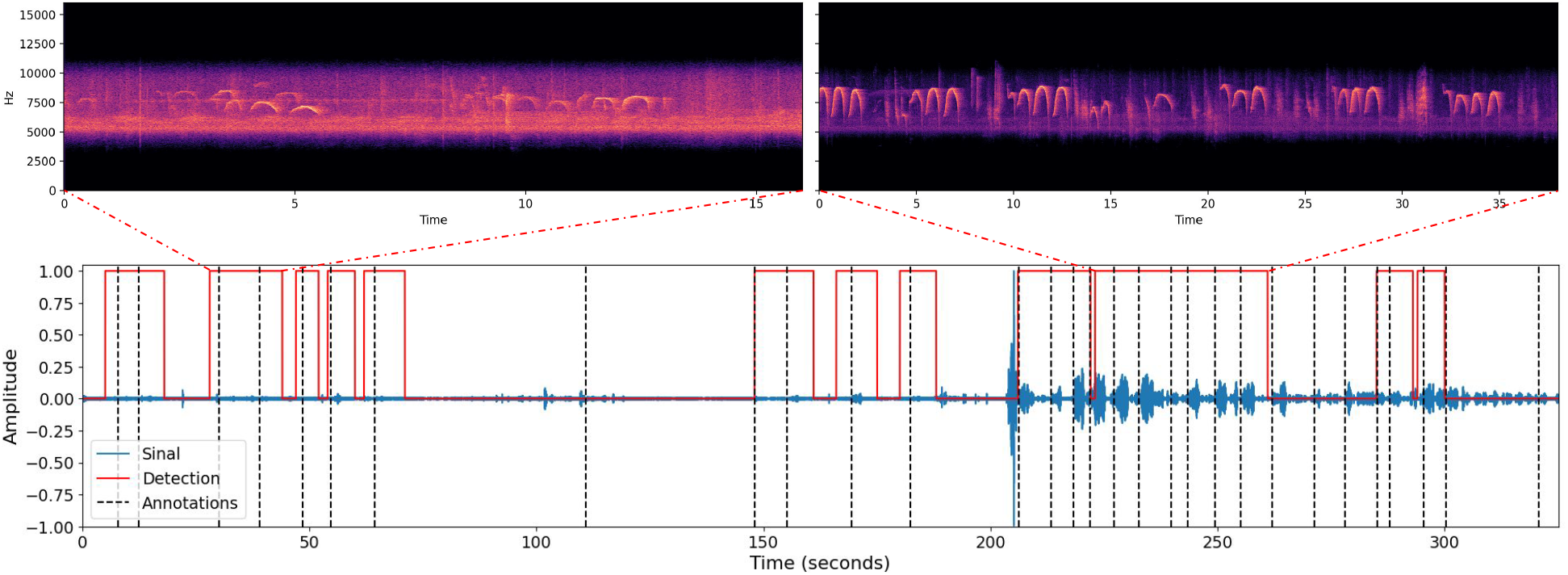
Detection timeline for a representative filtered recording of Mindú park. The waveform is shown with vertical dashed black lines marking the manually annotated onset of each *S. bicolor* call. Red lines denote the sequence of fixed-length time windows classified as inliers by the OCSVM, and spectrograms of the selected windows are displayed above. In this example, the detector yields no false positives but misses some calls (false negatives).

The red trace in Figure 7 marks the detected *S. bicolor* segments. Consecutive windows predicted with the same label are merged into continuous regions (solid red). We also show spectrograms for two contiguous intervals rich in calls, highlighting both the variability in call spectral shapes and changes in background noise, which is common in long recordings. In this example, we observe a false negative rate (FNR) of 69 % and a false positive rate (FPR) of 0.03 % (approximately zero).

To quantify detector quality, we matched model predictions to the human annotations using a simple temporal-overlap rule: a call is considered correctly detected if the human-annotated onset time falls within any predicted five-second positive window (equivalently, within ±2.5 s of that window’s center). This approximation is sufficient to compare and assess whether band-pass filtering prior to classification improves results.

Figure 8 shows ROC/AUC improvements when filtering is applied prior to feature extraction. These results reinforce that our method can be applied to new recordings from a different geographic region, with a different set of *S. bicolor* individuals and changing background conditions. Finally, the relatively small F1 values are driven by the lower recall (with high precision), indicating a detector that only labels *S. bicolor* when confidence is high, thereby avoiding false positives (Table 2).

**Table 2.** Detection performance on the Mindú Park example comparing OCSVM classification with and without band-pass filtering prior to feature extraction. Reported metrics are accuracy (Acc), precision (Pre), recall (Rec), F1-score (F1), and area under the ROC curve (AUC). Band-pass filtering improves AUC and F1 by raising true-positive rates while maintaining high precision.

| Band-pass | Acc | Pre | Rec | F1 | AUC |
| --- | --- | --- | --- | --- | --- |
| yes | 0.69 | 0.86 | 0.30 | 0.65 | 0.83 |
| no | 0.65 | 0.80 | 0.18 | 0.57 | 0.74 |

**Figure 8.**
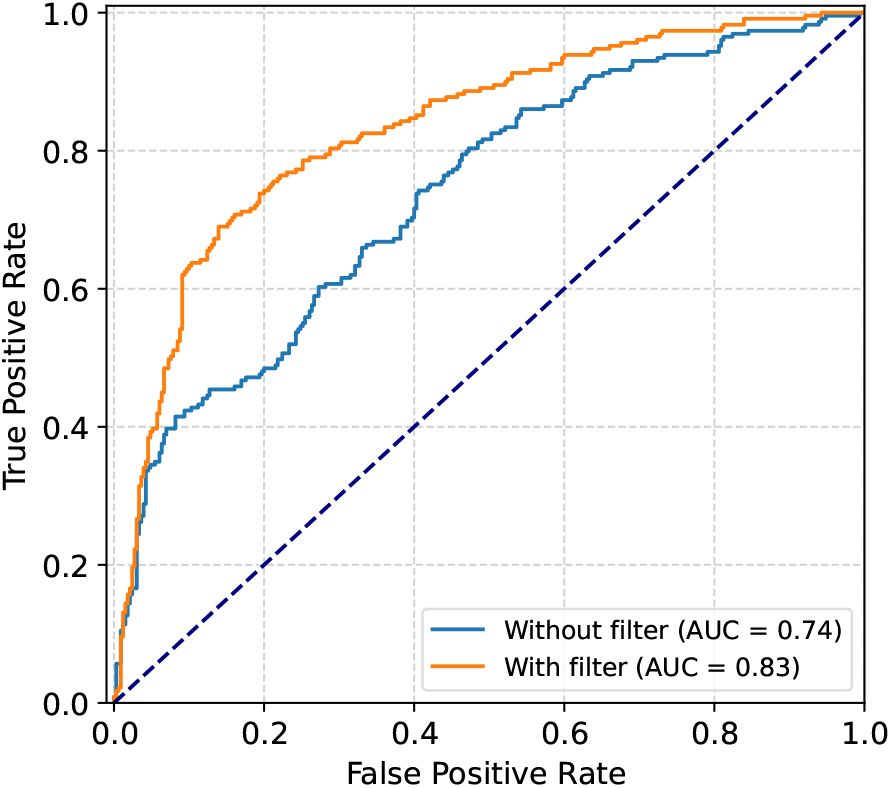
ROC curves for OCSVM-based detection computed on raw (unfiltered) audio and on band-pass–filtered audio. Band-pass filtering consistently improves performance across thresholds, yielding higher true positive rates and a larger area under the curve (AUC). The diagonal line denotes the random-classifier reference.

In the next section, we extend this cross-site evaluation by analyzing eight recordings from Mindú Park, geographically distant from the site used to train the OCSVM, to assess geographic transferability. Detection counts for these recordings are summarized in Table 3. We then show how to estimate detection probabilities at occupied site by plugging our detector into an occupancy model.

**Table 3.** Detection summary for Mindú Park recordings. Columns correspond to individual recording days *r*_1_, …, *r*_8_. *W* is the number of sliding windows extracted from each recording day; *D* is the number of windows detected as containing an *S. bicolor* call; and *y*_*t*_ indicates whether at least one detection occurred at visit *t*.

| | $r_1$ | $r_2$ | $r_3$ | $r_4$ | $r_5$ | $r_6$ | $r_7$ | $r_8$ |
| --- | --- | --- | --- | --- | --- | --- | --- | --- |
| $W$ | 757 | 149 | 109 | 309 | 1036 | 360 | 419 | 1320 |
| $D$ | 51 | 0 | 0 | 10 | 261 | 55 | 54 | 112 |
| $y_t$ | 1 | 0 | 0 | 1 | 1 | 1 | 1 | 1 |

## 7. Cross-site validation of the OCSVM detector with an occupancy model

Full timestamp annotation of all pied-tamarin vocalizations in PAM recordings, e.g., the timestamp labels of Section 6, is attention-intensive and time-consuming. A simpler alternative, namely, counting calls per unit time to estimate call density, still requires listening to entire files. When the objective is merely presence/absence at a site-visit, labeling can be further simplified by early stopping: listen only until the first confirmed vocalization, then stop. Despite its simplicity, these binary labels are sufficient for the occupancy framework presented by MacKenzie et al. (2003).

Our acoustic design follows the single-season occupancy framework of Royle and Link (2006); the key difference is that detections come from the OCSVM applied to acoustic recordings rather than from field observers.

Let *y*_*it*_ ∈ {0, 1} denote detection at site *i* on visit *t* (*t* = 1, …, *T*), and let *z*_*i*_ ∈ {0, 1} be the latent occupancy state with probability Ψ = Pr(*z*_*i*_ = 1). Assume conditional independence of visits given *z*_*i*_ and let 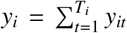 be the total number of detections at site *i* across *T*_*i*_ visits. Then, the likelihood of Ψ, *p*_11_, and *p*_10_ is

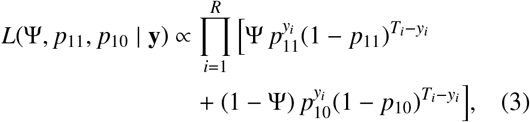

where *R* is the number of monitored sites, *y*_*i*_ is the total number of target detections at site *i* visited *T*_*i*_ times; and *p*_11_ = Pr(*y*_*i*_ = 1| *z*_*i*_ = 1) and *p*_10_ = Pr(*y*_*i*_ = 1 | *z*_*i*_ = 0) are the probabilities of detecting a pied tamarin when the site is occupied and of falsely detecting it when the site is unoccupied, respectively.

Because Ψ, *p*_11_, and *p*_10_ are unknown, they must be estimated jointly. The probabilities *p*_11_ and *p*_10_ depend on the detector’s error rates; with automated algorithms these are hard to specify *a priori* and typically require a hold-out dataset, which is unknown beforehand.

The Mindú Park dataset contains 38 recordings of varying lengths corresponding to eight monitoring days (Table 3), all labeled as target ‘1’ (pied tamarin present), so for these recordings we may take Ψ ≈ 1, i.e., treat *z*_*i*_ = 1. Under this assumption, the mixture component for unoccupied sites vanishes and the likelihood simplifies to the binomial form

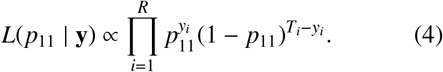

Note that with only occupied sites we can estimate *p*_11_, whereas *p*_10_ requires data from unoccupied sites.

In this set of experiments, we consider a single site (*R* = 1) visited on eight days (*T* = 8). The recordings come from a different *S. bicolor* group than the one used to train our classifier. Each recording has varying length and vocalization rate (Table 3). Because the numbers of windows *W* and detections *D*_*t*_ per visit can vary, we henceforth work with the binary indicator *y*_*t*_ = 1_(0,∞)_(*D*_*t*_), to mimic fieldwork procedures, which, when conducted in remote areas such as the Amazon rainforest, are typically organized by sampling days. Thus, any positive observation during a given day counts as a positive occupancy event for the species (*y*_*t*_ = 1). Under this reduction, Equation (4) simplifies to 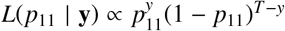, where 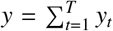. For computational convenience, we express the corresponding log-likelihood as

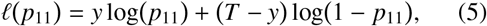

which yields the MLE 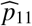 = *y*/*T* = 0.75 and the standard error

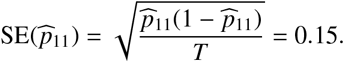

In our occupancy notation for occupied sites, the miss probability is *p*_01_ = *Pr*(*y*_*t*_ = 0 | *z* = 1) = 1 − *p*_11_. Thus, once we estimate 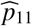, we immediately obtain 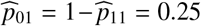. Appendix C shows the Maximum-Likelihood estimation (MLE) curve of *p*_01_. This helps characterize how well our detector can support a human expert when verifying PAM records.

## 8. Software package

To make the method accessible for conservation practitioners, we provide an open-source Python package, `sauim-detector`, distributed under the MIT license. The package can be conveniently installed via `pip` and runs locally. Given an audio file, the software applies the proposed pipeline (band-pass filtering, Perch embeddings, and OCSVM classification), and generates two outputs: (i) an Audacity-compatible label file (`.txt`) containing the detection timestamps, and (ii) a new `.wav` file representing a band-pass–filtered version of the original recording.

The label file can be directly imported into Audacity (`File` → `Import` → `Labels`), enabling analysts to visually inspect spectrograms and confirm *S. bicolor* calls, while the filtered audio provides a cleaner acoustic signal for playback or further analysis; cf. Figure 9. This design facilitates faster manual verification by focusing expert effort on candidate segments rather than entire recordings, thus reducing workload in large PAM datasets.

**Figure 9.**
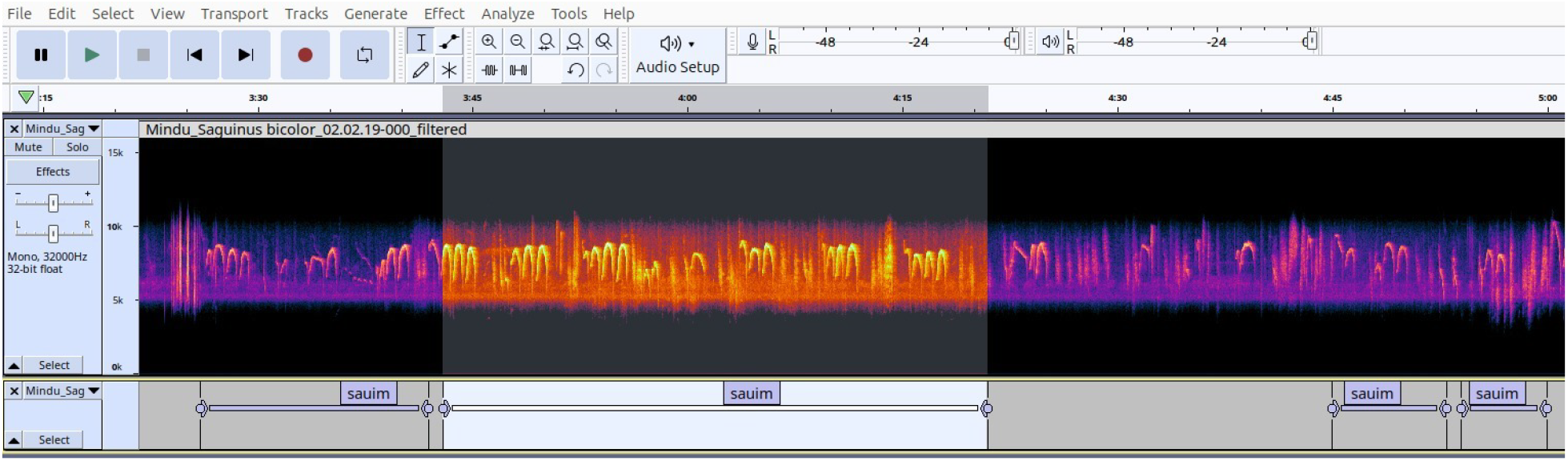
Example Audacity view after running `sauim-detector` and import the lables file. The spectrogram of the band-pass–filtered audio (top) and the imported label track highlighting candidate *S. bicolor* detections (bottom), enabling faster manual verification.

## 9. Discussion

### 9.1. Transfer learning and one-class detection in open acoustic environments

Across multiple non-target datasets that simulate open-set conditions, the proposed approach separated *S. bicolor* calls from other acoustic events and showed that a bird-oriented embedding model (Perch) can transfer effectively to a primate species. This result is consistent with a growing body of work showing that representations learned from large bioacoustic datasets can be transferred to species and acoustic domains that were not explicitly represented during model development. For example, Dufourq et al. (2022) demonstrated that transfer learning can substantially reduce the amount of labeled data required to develop PAM classifiers, achieving competitive performance with as few as 25 verified vocalizations using convolutional neural networks. Although their study did not evaluate Perch, their results support the broader premise that pretrained CNNs can be effectively adapted to bioacoustic detection tasks with limited labeled data.

More recently, few-shot transfer learning has been shown to improve species detection across heterogeneous landscapes while requiring only a small number of locally verified examples per class using Bird-Net (Jacuzzi and Olden, 2025). Our results extend this line of research in a different direction by using Perch as a general-purpose feature extractor rather than fine tuning its final classification layer. We then learn a one-class decision boundary using only examples of the target vocalization. This formulation is particularly attractive when the target species has few annotated examples and the negative class is difficult to enumerate. By not requiring an explicit representation of all possible non-target sounds during training, the approach avoids the need to anticipate every potential source of acoustic interference before deployment.

The trade-off, however, is that the learned decision boundary is determined by the variability represented in the positive training data. Consequently, the method should not be interpreted as capturing the full acoustic variability of *S. bicolor*; rather, it learns a decision boundary around the vocalizations represented in the available training data. This limitation is particularly relevant when deploying the detector across populations, recording conditions, or source-to-recorder distances that differ from those represented during training.

### 9.2. Specificity, sensitivity, and the role of automated pre-screening

The proposed method can scan long recordings using sliding windows, enabling both streaming detection and offline batch processing of large acoustic archives. This produces time-localized detections that can be directly incorporated into monitoring workflows.

In our cross-site detection evaluation, we obtained FPR = 0.03, FNR = 0.69, TPR = 0.31, and TNR = 0.97. These results indicate high specificity but limited sensitivity. This trade-off is important when interpreting the ecological usefulness of the proposed system. The detector should not currently be regarded as a replacement for expert inspection. Instead, its most immediate role is as a high-specificity pre-screening tool that can reduce the amount of audio requiring manual examination. The magnitude of this reduction will nevertheless depend on the prevalence of target vocalizations and the acoustic characteristics of the deployment environment.

For large PAM datasets, this distinction can have substantial practical consequences. The volume of continuously recorded audio can make exhaustive manual inspection impractical, and automated systems are increasingly being used to generate candidate detections for subsequent verification. Previous studies (Dufourq et al., 2022; Kath et al., 2024) have similarly emphasized the value of machine-learning pipelines for reducing manual inspection and annotation effort and making large acoustic datasets more tractable. Active-learning approaches further demonstrate that model predictions can be used to prioritize informative recordings and accelerate the creation of new training data (Kath et al., 2024).

### 9.3. Generalization across sites and the sensitivity– specificity trade-off

Our results indicate that Perch+OCSVM can transfer across different *S. bicolor* populations, while maintaining high specificity, despite possible variation in vocalization patterns. The cross-site experiment (Section 6) is particularly relevant because the training and evaluation recordings were obtained from different groups of individuals and from different acoustic environments. Thus, the evaluation introduces a distribution shift that is more representative of practical deployment than a random hold-out partition from the same recording conditions. The results suggest that the learned representation is sufficiently invariant to preserve specificity across these conditions, but not sufficiently invariant to preserve sensitivity.

Several factors may contribute to the limited sensitivity. Vocalizations may differ between individuals or groups, and previous behavioral work has shown that pied tamarins can modify their vocal behavior in response to anthropogenic noise (Sobroza et al., 2024). In addition, acoustic attenuation increases with distance between the animal and the recorder, particularly at higher frequencies, reducing signal-to-noise ratio and potentially causing genuine calls to become less similar to the target examples represented in the training set. These effects are especially relevant for PAM because the distance and orientation of a vocalizing animal relative to a fixed recorder are generally unknown.

The band-pass filter provides an example of how ecological knowledge can partially compensate for these sources of variation. Because *S. bicolor* vocalizations concentrate much of their energy in the 5–10 kHz range, filtering suppresses portions of the soundscape that are less informative for the target species. The improvement observed after filtering is consistent with the broader principle that domain knowledge remains valuable even when using deep pretrained representations.

The possibility of confusion with closely related taxa is another important consideration. In particular, *Saguinus midas* occurs in part of the geographic range of *S. bicolor*, and convergent acoustic characteristics between sympatric tamarin species have been documented (Sobroza et al., 2021). The negative datasets used in the present study do not provide a systematic test against *S. midas*. Consequently, the current results should not yet be interpreted as demonstrating species-level discrimination under all sympatric conditions. Evaluation against sympatric and acoustically similar species is therefore an important requirement for assessing the robustness of the detector under realistic deployment conditions.

### 9.4. From acoustic detections to ecological inference

The results shown in Section 6 should be interpreted with some caution because onset timestamps may be slightly misaligned with the detector’s analysis windows, which can affect event-matching metrics. In addition, we observed that vocalizations with two or fewer bouts tend to increase FNR and can degrade PAM performance.

In large PAM datasets spanning many hours of audio, it is common to pre-screen recordings with an automated detector to generate candidate timestamps. A human expert then reviews only these candidates to confirm that each detection corresponds to the target species (e.g., pied tamarin), thereby reducing manual time and effort. For this verification workflow, high sensitivity is desirable because missed vocalizations cannot be recovered by reviewing only the candidate events. However, the relatively high FNR observed in our cross-site evaluation limits the use of the current detector as a standalone pre-screening mechanism when the objective is to minimize missed vocalizations. In this context, the software package presented in Section 8 has the potential to substantially reduce the human effort required to inspect new field survey recordings for the species.

Building on these detections, we link the detector outputs to a simplified single-site occupancy framework (Section 7), motivated by the broader idea that occupancy-style approaches can be useful for interpreting machine-learning detections under imperfect detection. Assuming Mindú Park is occupied (Ψ = 1), the MLE has the closed form 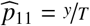, where *y* is the number of visits with at least one detection across *T* visits; the corresponding visit-level non-detection probability is *p*_01_ = 1 − *p*_11_ = 0.25. This quantity should not be confused with the event-level FNR = 0.69 reported above. The two quantities describe different levels of the detection process: the event-level FNR concerns individual acoustic events, whereas *p*_01_ concerns the probability that an entire visit produces no detection. Because each visit contains multiple opportunities for detection, its probability of producing at least one detection can be substantially higher than the probability of detecting any particular vocalization.

Although simple, this example illustrates how detector outputs can be incorporated into an occupancy-oriented analysis. However, the primary goal of the present study is not to propose a complete ecological occupancy framework, but rather to explore whether an occupancy-style formulation can provide an interpretable summary of the behavior of a one-class bioacoustic detector under imperfect detection conditions.

Recent PAM studies have similarly emphasized the importance of integrating automated acoustic detection with ecological models rather than treating classification accuracy as the final monitoring objective. For example, Wood et al. (2023b) combined BirdNET-based acoustic detection with occupancy modeling to monitor an endangered primate, illustrating how automated detections can be translated into ecologically interpretable measures. In contrast, our formulation uses one-class learning, allowing the detector to be trained using only target-species vocalizations, without requiring an explicit representation of the potentially unbounded set of non-target sounds.

Under manual inspection, false positives impose an additional inspection cost, whereas false negatives can lead to an ecological inference error if missed vocalizations are interpreted as evidence of absence. Because multiple monitoring visits provide repeated opportunities for detection, the probability of detecting the species can increase with repeated sampling, partly offsetting this risk. Nevertheless, an important caveat applies: the Mindú Park recordings were collected manually and deliberately close to the acoustic source (near *S. bicolor*); in PAM programs with fixed-position recorders, the source is not always close to the microphone, which can reduce SNR and limit the transferability of these detection-probability estimates.

From a conservation perspective, the main contribution of the proposed system is not the replacement of conventional field surveys, but their augmentation. By transforming long recordings into a smaller set of candidate events, the detector can help make PAM more scalable and reduce the effort required for expert inspection. This is particularly relevant for *S. bicolor*, for which acoustic monitoring could complement conventional field-based surveys in fragmented and difficult-to-access habitats.

## 10. Conclusions

We presented an open-set bioacoustic detection pipeline for the critically endangered pied tamarin (*Saguinus bicolor*), combining a domain-informed band-pass filter, pretrained Perch embeddings, and a One-Class Support Vector Machine (OCSVM) optimized through a Leave-One-Out cross-validation strategy. The main contribution is the demonstration that a pretrained bioacoustic representation originally developed for large-scale bird classification can be effectively repurposed for a primate species using a relatively small number of target examples, without requiring an explicit enumeration of the negative acoustic class.

The detector achieved high specificity in cross-site recordings, although event-level sensitivity remained limited. These results support the use of the proposed system primarily as a candidate-generation and prescreening tool to reduce the amount of audio requiring expert inspection, rather than as a standalone mechanism for establishing species absence or estimating call abundance. The integration of automated detections with an occupancy-oriented formulation further illustrates how imperfect acoustic detections can be connected to ecological inference.

Future work should evaluate the approach across multiple sites, including different individuals, environmental conditions, source-to-recorder distances, and sympatric species such as *Saguinus midas*. Alternative pretrained embedding models, modest fine-tuning on *S. bicolor* vocalizations, and active or semi-supervised learning could further improve sensitivity and robustness to novel acoustic environments.

Finally, multi-season studies incorporating ecological covariates such as seasonal variation and habitat characteristics could further establish the ecological value of automated detections. Together with the publicly released data and software, these developments can contribute to more scalable and reproducible acoustic monitoring of *S. bicolor* and other threatened species in complex tropical soundscapes.

## Acknowledgements

This work was supported by the Coordenação de Aperfeiçoamento de Pessoal de Nível Superior – Brazil (CAPES-PROEX), Financing Code 001, and by the Fundação de Amparo à Pesquisa do Estado do Amazonas (FAPEAM), through the PDPG/CAPES and POSGRAD 2026/2027 projects. We also thank FAPEAM for financial support through the project “Diferentes Abordagens Computacionais para Monitoramento Ecoacústico Autônomo da Região Amazônica”, Public Call No. 013/2022 – Produtividade em CT&I. JGC thanks OpenAI for its support through the partnership established with OpenAI representative Nicolas Robinson Andrade as part of the AmazonGPT project. TVS thanks The Rufford Foundation (24762-1 and 37915-2), the National Geographic Society (EC-419R-18), and the International Primatological Society for funding fieldwork. ACF thanks MBIE/NASA Catalyst for its support through the project “Monitoring Vegetation-Geothermal Interactions from Space and Airborne Platforms”, project number E14490.

## Data Availability

The data and source code supporting the findings of this study are publicly available at https://github.com/juancolonna/Sauim/tree/main. The repository contains the audio recordings, manual annotations, and other data used in the experiments, as well as the source code required to reproduce the analyses and results presented in this study.

## Appendix A. Species

### Bird dataset species composition

*Cacicus haemorrhous, Cercomacroides nigrescens, Chiroxiphia pareola, Chrysomus icterocephalus, Glaucidium hardyi, Graydidascalus brachyurus, Hemitriccus minor pallens, Herpsilochmus dorsimaculatus, Hypocnemis flavescens, Ibycter americanus, Leistes militaris, Myiopagis gaimardii, Myrmoborus lugubris stictopterus, Neopelma chrysocephalum, Pharomachrus pavoninus, Pionus fuscus, Pseudopipra pipra, Ramphocelus carbo, Rupicola rupicola, Schiffornis olivacea, Sublegatus obscurior, Thamnophilus amazonicus cinereiceps, Thamnophilus murinus, Todirostrum pictum, Tyrannus melancholicus, Zimmerius acer*

### Anuran dataset species composition

*Adenomera hylaedactyla, Hyla minuta, Adenomera andreae, Ameerega trivittata, Osteocephalus oophagus, Rhinella granulosa, Scinax ruber, Hypsiboas cinerascens, Hypsiboas cordobae, Leptodactylus fuscus*

## Appendix B. Confusion matrices

**Figure B.10.**
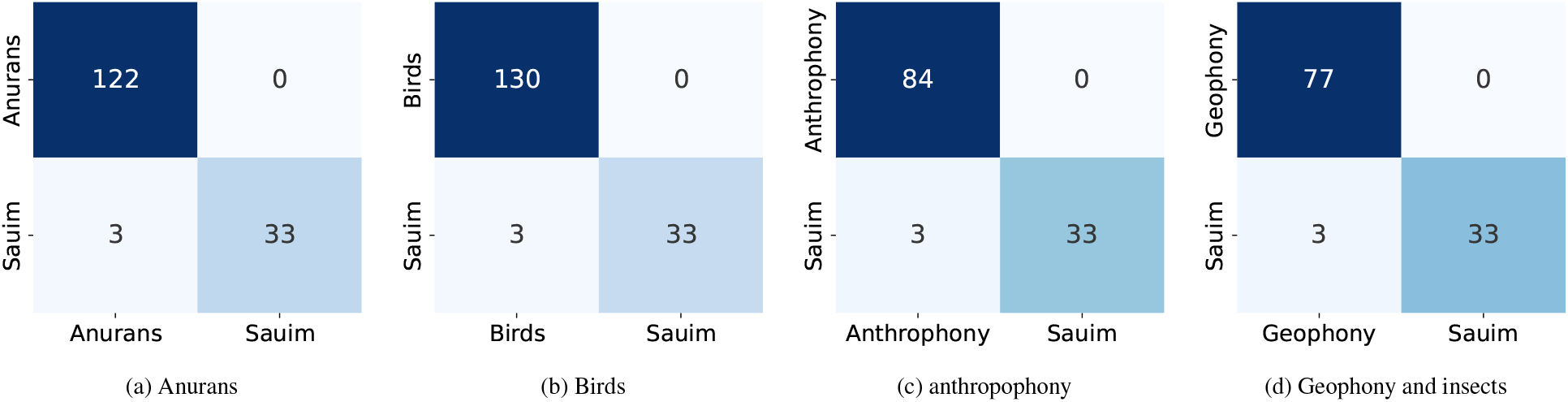
Confusion matrices for the OCSVM with band-pass filtering. Each matrix reports TP, FP, TN, and FN counts on the hold-out partition using the default decision threshold.

## Appendix C Maximum likelihood estimation

**Figure C.11.**
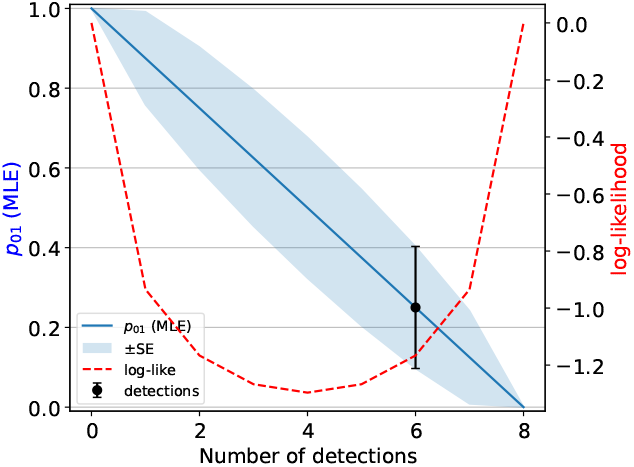
MLE of 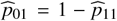 as a function of the number of detections *k*. The blue curve shows 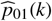 with a shaded ±1 standard error band. The black point highlights the estimate at the observed detections *k* = Σ_*i*_ **1**{*y*_*i*_ > 0}, with its error bar. The red dashed line (right axis) shows the maximized log-likelihood max_*p*11_ log *L*(*k* | *p*_11_) as a function of *k*.

## Appendix D Cluster of embeddings vectors

**Figure D.12.**
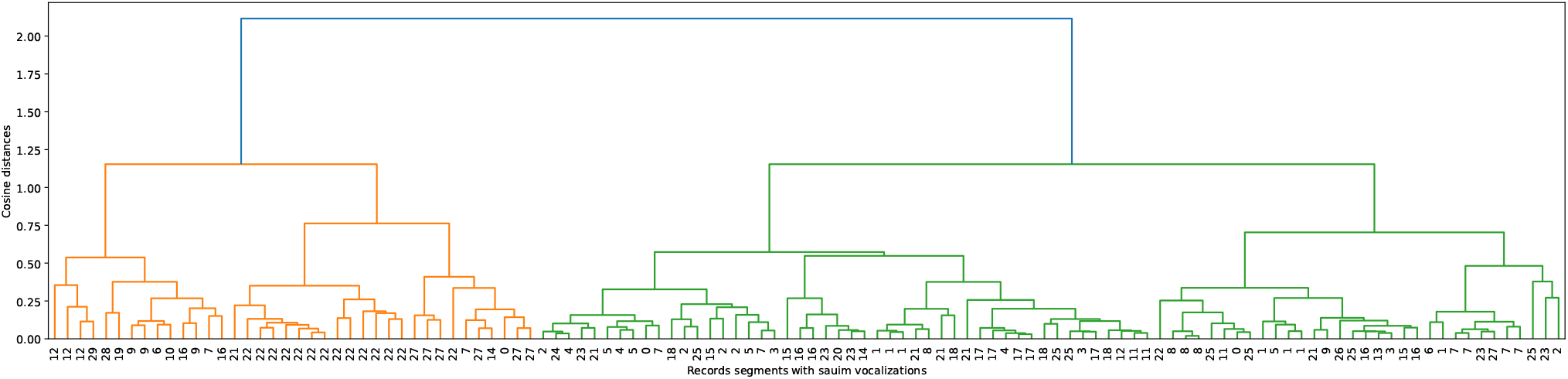
Hierarchical clustering dendrogram of target-class *S. bicolor* embeddings (cosine distance, average linkage). Each leaf is labeled with the integer ID of the recording from which the call segment was extracted. Segments from the same recording tend to cluster together, likely reflecting the same individual and shared background noise.

## Footnotes

1 https://xeno-canto.org/

2 https://docs.scipy.org/doc/scipy/reference/generated/scipy.signal.sosfiltfilt.html

